# Intron-containing RNA from the HIV-1 provirus activates type I interferon and inflammatory cytokines

**DOI:** 10.1101/128447

**Authors:** Sean Matthew McCauley, Kyusik Kim, Anetta Nowosielska, Ann Dauphin, Leonid Yurkovetskiy, William Edward Diehl, Jeremy Luban

## Abstract

HIV-1-infected people who take drugs that suppress viremia to undetectable levels are protected from developing AIDS. Nonetheless, these individuals have chronic inflammation associated with heightened risk of cardiovascular pathology. HIV-1 establishes proviruses in long-lived CD4^+^ memory T cells, and perhaps other cell types, that preclude elimination of the virus even after years of continuous antiviral therapy. Though the majority of proviruses that persist during antiviral therapy are defective for production of infectious virions, many are expressed, raising the possibility that the HIV-1provirus or its transcripts contribute to ongoing inflammation. Here we found that the HIV-1 provirus activated innate immune signaling in isolated dendritic cells, macrophages, and CD4^+^ T cells. Immune activation required transcription from the HIV-1 provirus and expression of CRM1-dependent, Rev-dependent, RRE-containing, unspliced HIV-1 RNA. If *rev* was provided *in trans*, all HIV-1 coding sequences were dispensable for activation except those *cis*-acting sequences required for replication or splicing. These results indicate that the complex, post-transcriptional regulation intrinsic to HIV-1 RNA is detected by the innate immune system as a danger signal, and that drugs which disrupt HIV-1 transcription or HIV-1 RNA metabolism would add qualitative benefit to current antiviral drug regimens.

## INTRODUCTION

Drugs that block activity of the three HIV-1 enzymes-reverse transcriptase, integrase, and protease-potently suppress HIV-1 viremia and protect infected people from developing AIDS (Günthard et al., 2016). Despite effective antiviral therapy, many patients have systemic inflammation associated with increased risk of cardiovascular pathology (Brenchley et al., 2006; Freiberg et al., 2013; Sinha et al., 2016). Plausible explanations for this inflammation include ongoing T cell dysfunction, disruption of intestinal epithelium integrity, lymphatic tissue fibrosis, antiviral drug toxicity, and co-morbid infections such as cytomegalovirus (Brenchley et al., 2006; Hunt et al., 2011; Sinha et al., 2016).

Inflammation may also be maintained by HIV-1 itself. HIV-1 is not eliminated from the body after years of continuous suppressive therapy and viremia inevitably rebounds upon drug cessation (Davey et al., 1999). This is because HIV-1 establishes a provirus in long-lived CD4^+^ memory T cells and perhaps other cell types that include tissue-resident myeloid cells (Jiang et al., 2015; Kandathil et al., 2016; Siliciano et al., 2003). Proviruses are obligate replication intermediates that result from the integration of retroviral cDNA into chromosomal DNA to become permanent, heritable genetic elements in infected cells (Engelman and Singh, 2018). Though the vast majority of proviruses that persist in the presence of antiviral therapy are defective for the production of infectious virus (Bruner et al., 2016), 2-18% express HIV-1 RNA (Wiegand et al., 2017). Here we assessed HIV-1 proviruses and their gene products for the ability to contribute to chronic inflammation.

## RESULTS

To determine if HIV-1 proviruses activate innate immune signaling, human blood cells were transduced with single-cycle vectors, either a full-length, single-cycle HIV-1 clone with a frameshift in *env* and eGFP in place of *nef* (HIV-1-GFP) (He et al., 1997), or a minimal 3-part lentivector encoding GFP (Fig. 1a and Table 1) (Pertel et al., 2011). Monocyte derived dendritic cells (DCs) were challenged initially since HIV-1 transduction of these specialized antigen-presenting cells activates innate immune signaling (Berg et al., 2012; Gao et al., 2013; Landau, 2014; Manel et al., 2010; Rasaiyaah et al., 2013). To increase the efficiency of provirus establishment, vectors were pseudotyped with the vesicular stomatitis virus glycoprotein (VSV G) and delivered concurrently with virus-like particles (VLPs) bearing SIV_MAC_251 Vpx (Fig. 1b) (Goujon et al., 2006; Pertel et al., 2011). Transduction efficiency, as determined by flow cytometry for GFP-positive cells, was 30-60% (Fig. 1b), depending on the blood donor.

**Table 1.**
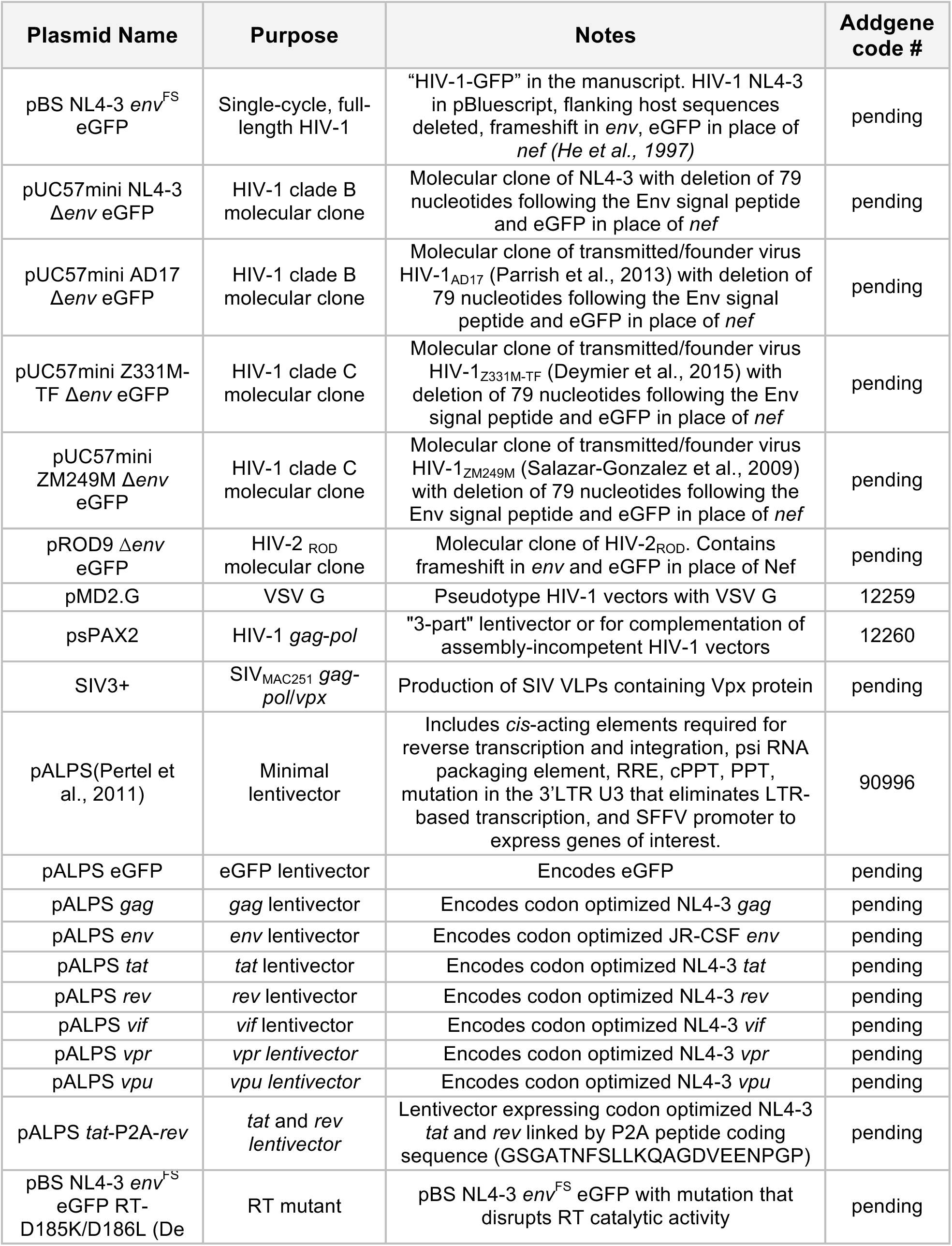
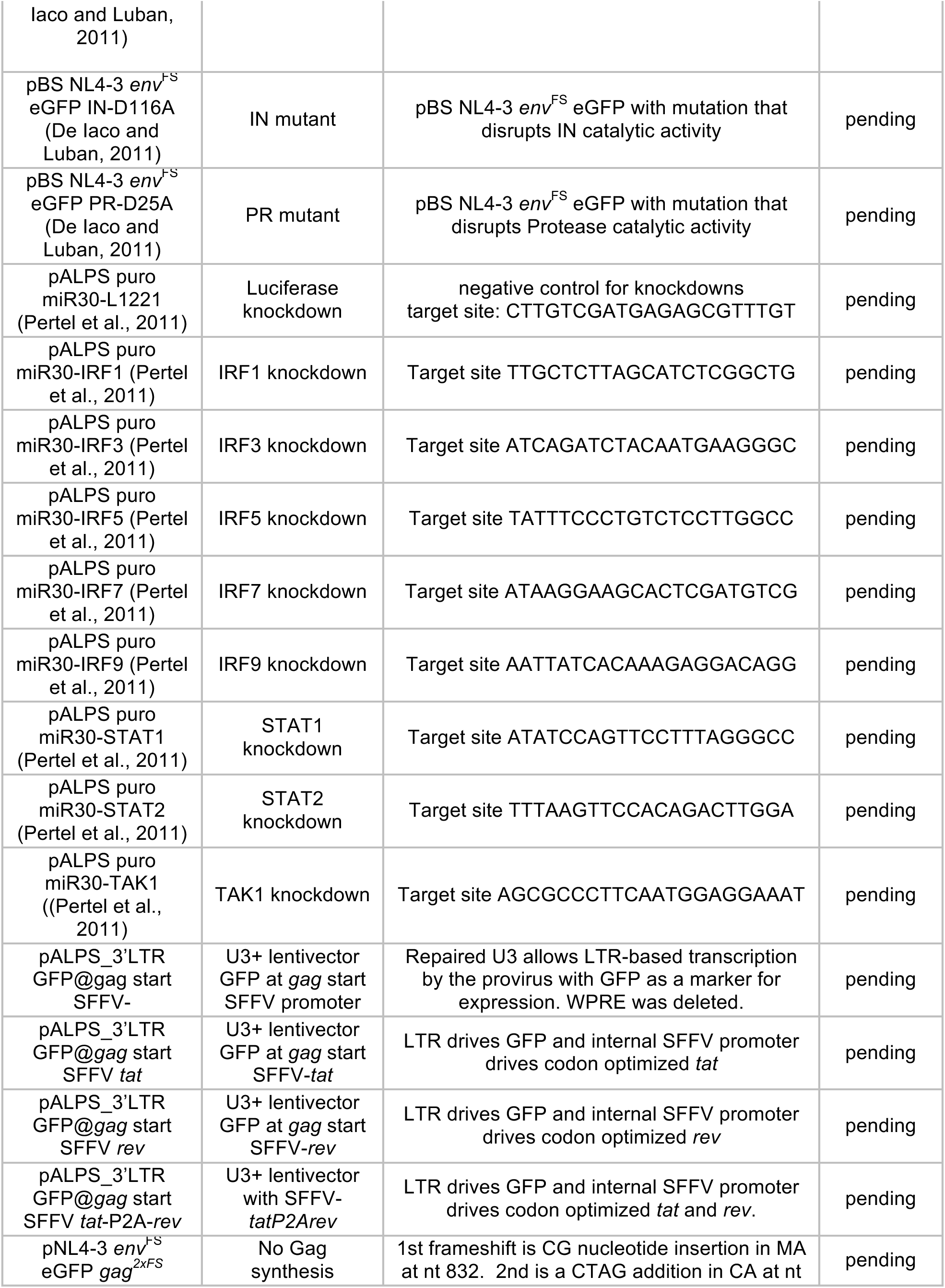
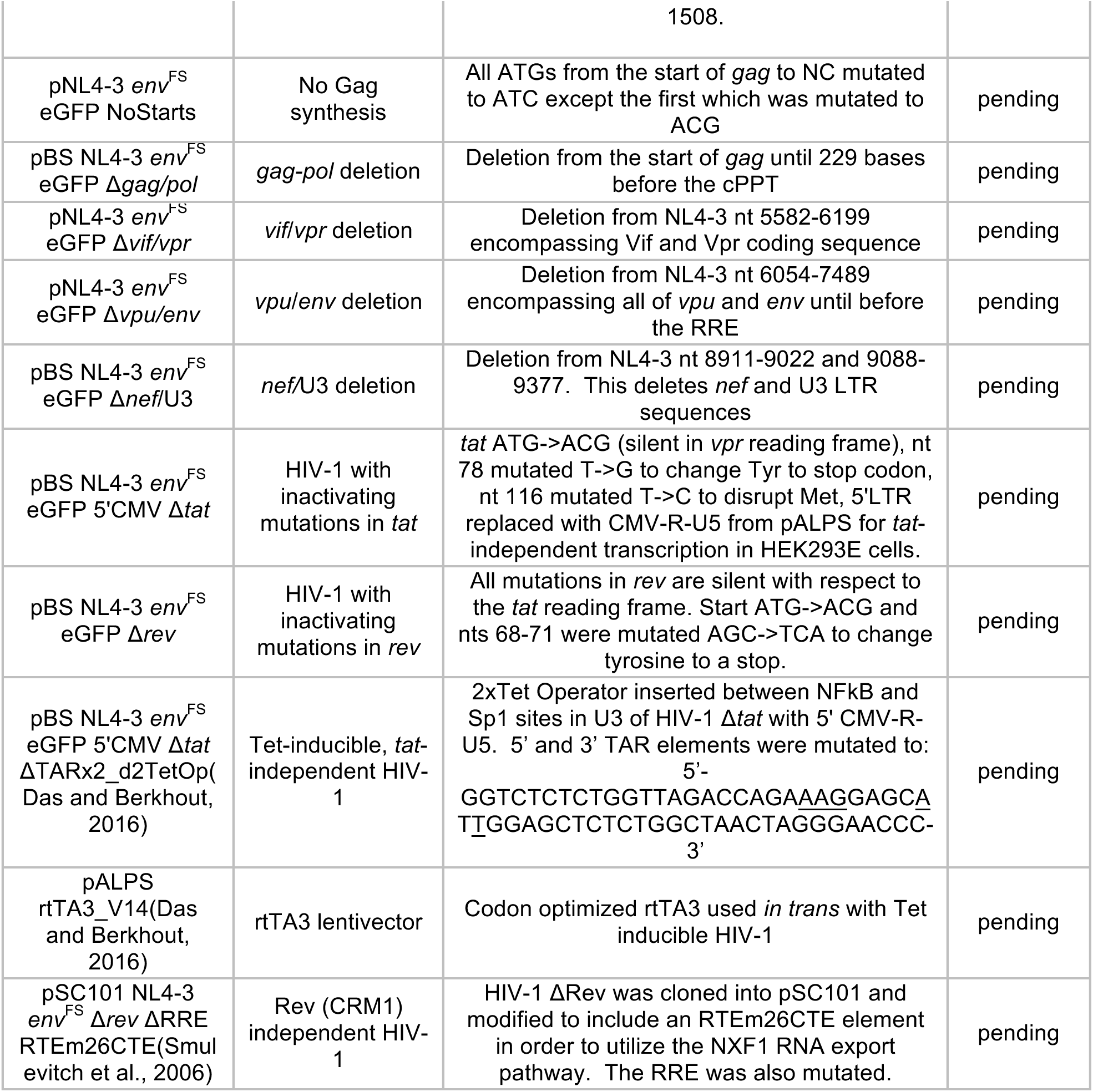
Plasmids used in this study.

**Figure 1.**
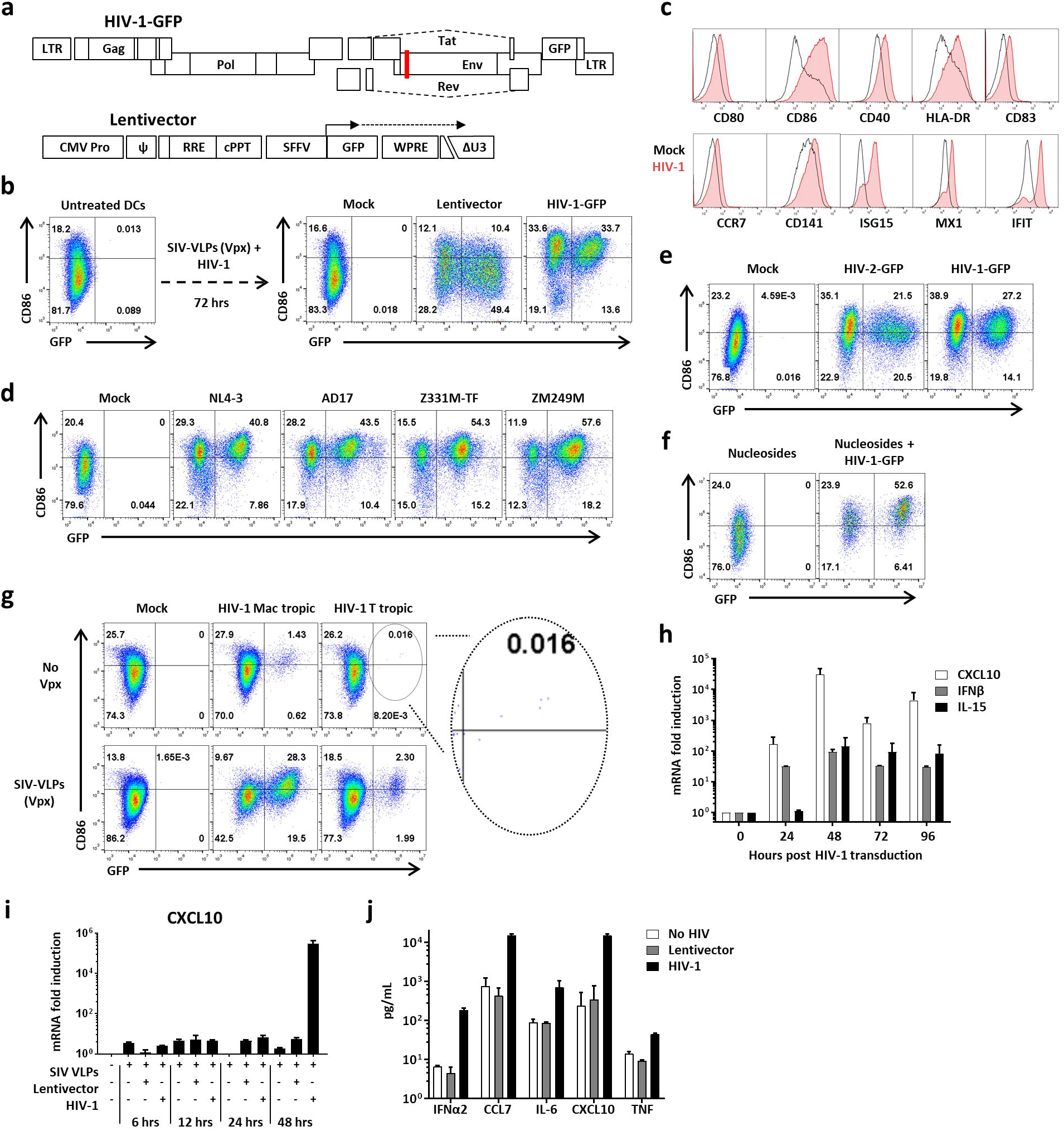
HIV-1 transduction matures DCs. **a,** Schematic of HIV-1-GFP, with frameshift in *env* (red line) and *gfp* in place of *nef(He et al., 1997)*, and of the minimal lentivector, with self-inactivating ΔU3 LTR(Zufferey et al., 1998) and *gfp* driven by the SFFV promoter (Pertel et al., 2011). Unless indicated otherwise, vectors were pseudotyped with VSV G and cells were co-transduced with SIV_MAC_251 VLPs bearing Vpx. **b**, Flow cytometry of DCs for GFP and CD86, after treatment as indicated. **c,** Flow cytometry histograms for the indicated markers 72 hrs after DC transduction with HIV-1 (red) or mock (black). **d,** Flow cytometry of DCs for GFP and CD86 after transduction with single-cycle clones, HIV-1_NL4-3_, HIV-1_AD17_, HIV-1_Z331M-TF_, or HIV-1_ZM249M_. **e,** Transduction of DCs with HIV-2_ROD_-GFP, single-cycle vector. **f,** DC transduction with HIV-1-GFP in the absence of Vpx and the presence of 2 mM nucleosides. **g,** 12 day spreading infection on DCs, with macrophage-tropic or T cell-tropic, replication-competent HIV-1, with or without SIV VLPs. **h,** qRT-PCR quantitation of *CXCL10* (black), *IFNB1* (gray), or *IL15* (white) mRNAs from DCs transduced with HIV-1-GFP. **i,** qRT-PCR quantitation of *CXCL10* mRNA in DCs transduced with either HIV-1-GFP or minimal lentivector, assessed at the indicated times post-transduction. **j,** Cytokines in DC supernatant as assessed by luminex, 72 hrs after transduction with HIV-1-GFP (black) or minimal lentivector (gray). Shown are blood donor data representative of n=12 (**b**), n=4 (**c, d, e, f, h, i, j**), or n=8 (**g**). To determine significance, the MFI of all live cells for each sample was calculated as fold-change versus control. The exception being (**g**) where the MFI of only GFP+ cells was compared. When data from each donor replicate within a experiment was combined, the difference in MFI for all experimental vs control conditions was significant in all cases, p<0.0001; one-way ANOVA, Dunnett’s post-test. qRT-PCR and Luminex data were mean +/-SD, p < 0.0001; two-way ANOVA, Dunnett’s post-test.

DCs matured in response to HIV-1-GFP transduction, as indicated by increased mean fluorescence intensity of co-stimulatory or activation molecules, including HLA-DR, CD80, CD86, CD40, CD83, CCR7, CD141, ISG15, MX1, and IFIT (Sousa, 2006) (Fig. 1b, c). Maturation was evident among both GFP positive and negative cells (Fig. 1b), the latter resulting from activation *in trans* by type 1 IFN as several others have shown (Manel et al., 2010; Rasaiyaah et al., 2013). Identical results were obtained with full-length, single-cycle vectors generated from primary, transmitted/founder clones that were derived by single genome sequencing, HIV-1_AD17_ (Parrish et al., 2013), HIV-1_Z331M-TF_ (Deymier et al., 2015), and HIV-1_ZM249M_ (Salazar-Gonzalez et al., 2009), the first virus being clade B, the other two clade C (Fig. 1d). A single cycle HIV-2 vector also induced maturation, indicating that this innate response was not unique to HIV-1 (Fig. 1e).

DCs matured when HIV-1-GFP transduction efficiency was augmented with nucleosides (Reinhard et al., 2014), rather than with SIV VLPs, indicating that Vpx was not required for maturation (Fig. 1f). DCs were then challenged with replication-competent HIV-1 bearing CCR5-tropic Env, either T cell-tropic or macrophage-tropic (Granelli-Piperno et al., 1998), with or without Vpx-VLPs (Fig. 1g). The percent of cells transduced by vector bearing either Env increased with Vpx, though DC maturation was observed under all conditions, even among the very few DCs transduced by T cell-tropic *env* (see inset of Fig. 1g). These results indicate that neither VSV G, nor Vpx, nor high-titer infection, was required for DC maturation.

In response to transduction with HIV-1-GFP, steady-state *CXCL10*, *IFNB1*, and *IL15* mRNAs reached maximum levels at 48 hrs, increasing 31,000-, 92-, and 140-fold relative to mock-treated cells, respectively (Fig. 1h, i). Correspondingly, IFNα2, CCL7, IL-6, CXCL10, and TNFα proteins accumulated in the supernatant (Fig. 1j). In contrast to the results with HIV-1-GFP, there were no signs of maturation after transduction with the 3-part minimal lentivector (Fig. 1b, h, j).

To determine if early stages in the HIV-1 replication cycle were necessary for maturation, reverse transcription was inhibited by nevirapine (NVP) or the HIV-1 RT mutant D185K/D186L, and integration was inhibited with raltegravir or the HIV-1 IN mutant D116A (Tables 1 and 2), as previously described (De Iaco and Luban, 2011). Each of these four conditions abrogated maturation, as indicated by cell surface CD86 (Fig. 2a) and steady-state *CXCL10* mRNA (Fig. 2b). When integration was inhibited, *CXCL10* mRNA increased in response to challenge with HIV-1-GFP, but levels were nearly 1,000 times lower than when integration was not blocked (Fig. 2b).

**Table 2.** Drugs and reagents.

| **Drug** | **Action** | **Source** | **Working concentration** | **HIV-1 DC maturation** |
| --- | --- | --- | --- | --- |
| Doxycycline | rtTA3 activator | Sigma (D9891) | 10-1000 ng/mL | - |
| cGAMP | STING activator | Invivogen (tlrl-nacga23) | 25 µg/mL | - |
| PHA-P | T cell mitogen | Sigma (L1668) | 5 µg/mL | - |
| 2'deoxyguanosine monohydrate | For nucleoside assisted transductions | Sigma (D0901) | 2 mM | - |
| 2' deoxythymidine | For nucleoside assisted transductions | Sigma (T1895) | 2 mM | - |
| 2'deoxyadenosine monohydrate | For nucleoside assisted transductions | Sigma (D8668) | 2 mM | - |
| 2'deoxycytidine hydrochloride | For nucleoside assisted transductions | Sigma (D0776) | 2 mM | - |
| Sendai Virus (SeV) Cantell strain | Challenge virus | Charles River Labs (VR-907) | 200 HA units/mL | - |
| Nevirapine | Reverse transcriptase inhibitor | NIH AIDS reagent program (4666) | 5 µM | Inhibits |
| Raltegravir | Integrase inhibitor | NIH AIDS reagent program (11680) | 10 µM | Inhibits |
| Leptomycin | CRM1 inhibitor | Invivogen (tlrl-lep) | 25 nM | Inhibits |
| Cyclosporin A | Cyclophilin A inhibitor | Sigma (30024) | 5 µM | No effect |
| GW-5075 | c-Raf | Sigma (G6416) | 1, 5, 25 µM | No effect |
| BAY11-7082 | IκB-α Inhibitor | Invivogen (tlrl-b82) | 1, 2.5, 10 µM | No effect |
| U0126 | MEK1 and MEK2 Inhibitor | Invivogen (tlrl-u0126) | 10, 25, 50 µM | No effect |
| SB203580 | p38 MAP Kinase Inhibitor | Invivogen (tlrl-sb20) | 1, 2.5, 10 µM | No effect |
| MCC950 | NLRP3-inflammasome inhibitor | Invivogen (inh-mcc) | 1, 2.5, 10 µM | No effect |
| SP600125 | JNK Inhibitor | Invivogen (tlrl-sp60) | 10, 25, 100 µM | No effect |
| Z-VAD-FMK | Pan-Caspase Inhibitor | Invivogen (tlrl-vad) | 1, 5, 20 μM | No effect |
| NQDI-1 | ASK1 inhibitor | Sigma (SML0185) | 1, 10, 100 μM | No effect |
| ISRIB | eIF2a phosphorylation inhibitor | Sigma (SML0843) | 1, 10, 100 μM | No effect |
| VX-765 | Caspase 1 inhibitor | Invivogen (inh-vx765i) | 1, 10, 100 μM | No effect |
| Dexamethasone | NF-kB and MAPK inhibitor | Invivogen (tlrl-dex) | 10, 100, 1000 nM | No effect |
| Chloroquine | inhibitor of endosomal acidification | Invivogen (tlrl-chq) | 1, 10, 100 μM | No effect |
| Amlexanox | TBK1/IKKε inhibitor | Invivogen (inh-amx) | 1, 10, 100 μg/mL | No effect |
| BX795 | TBK1/IKKε inhibitor | Invivogen (tlrl-bx7) | 0.5, 1, 2 μM | No effect |
| NG25 trihydrochloride | TAK1 & LYN, MAP4K2 and Abl inhibitor | Sigma (SML1332) | 50, 100, 500, 1000 nM | No effect |
| 5Z-7-Oxozeaenol | TAK1 & MAP4K2 inhibitor | Sigma (O9890) | 50, 100, 500, 1000 nM | No effect |
| C16 | PKR inhibitor | Sigma (I9785) | 1 μM | No effect |
| 2AP | PKR inhibitor | Invivogen (tlrl-apr) | 5 μM | No effect |

**Figure 2.**
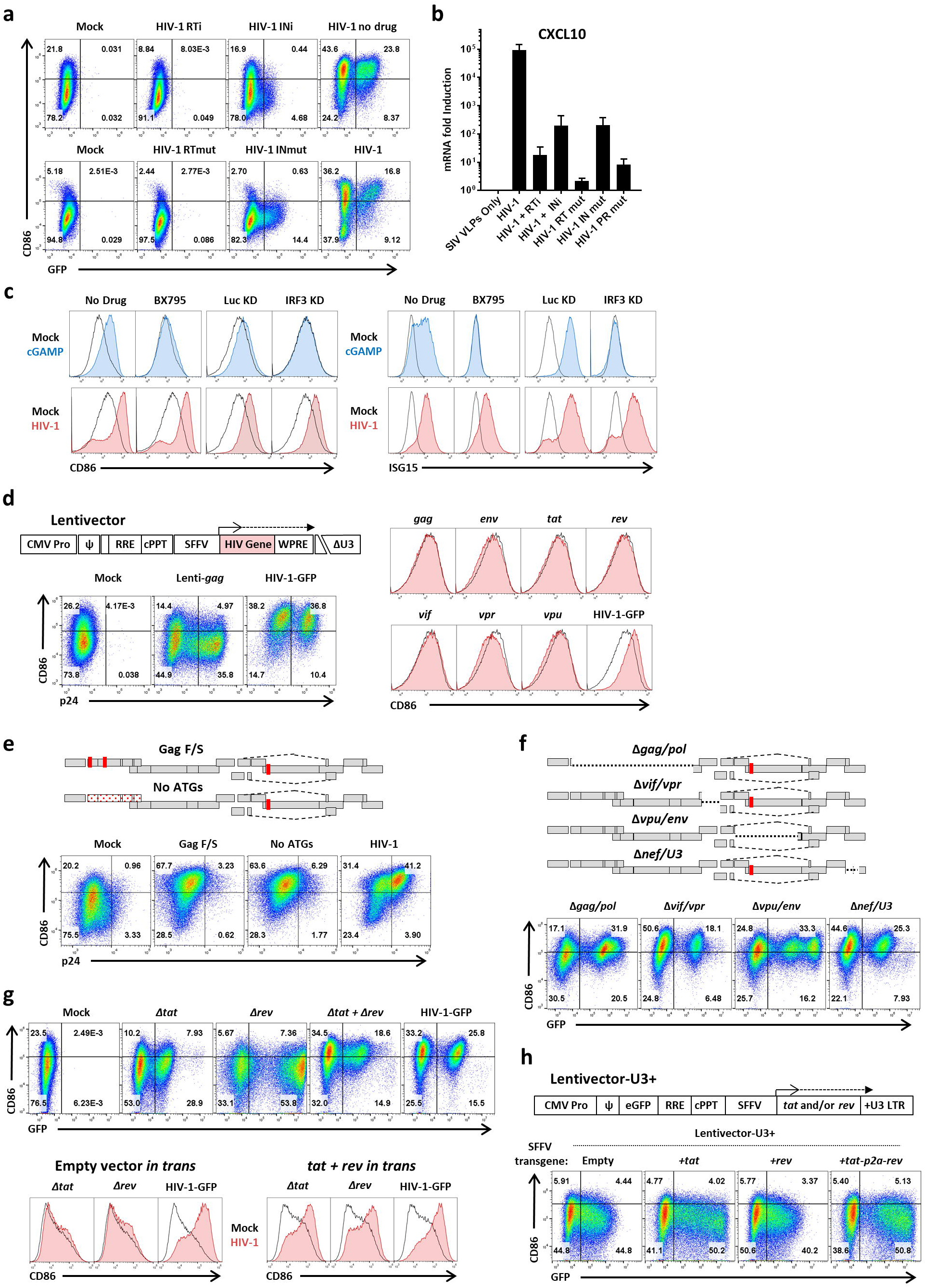
Native HIV-1 RNA regulation is necessary for DC maturation, but LTR-driven transcription is insufficient and coding sequences are not necessary. **a,** Assessment of GFP and CD86 by flow cytometry following transduction with, top, HIV-1-GFP in the presence of 5 µM nevirapine (RTi), 10 µM raltegravir (INi), or no drug, and, bottom, HIV-1-GFP bearing mutant RT-D185K/D186L (RTmut) or mutant IN-D116A (INmut). **b**, qRT-PCR quantitation of *CXCL10* mRNA from the same DCs as in (**a**). **c,** DCs treated with 1 µM of the TBK1 inhibitor BX795, or expressing shRNAs targeting either IRF3 or luciferase control (Pertel et al., 2011), were challenged with 25 µg/mL cGAMP or HIV-1-GFP and assayed by flow cytometry for CD86 and ISG15. **d,** Flow cytometry of DCs after transduction with minimal lentivectors expressing codon optimized HIV-1 genes; **e**, HIV-1-GFP in which translation was disrupted by two frameshifts in *gag* or by mutation of the first 14 AUGs in *gag*; **f,** HIV-1-GFP bearing deletion mutations encompassing *gag*/*pol*, *vif*/*vpr*, *vpu*/*env*, or *nef*/U3-LTR; **g,** HIV-1-GFP bearing mutations in *tat* or *rev*, co-transduced with both mutants, or co-transduced with minimal vector expressing *tat* and *rev in trans*; or **h,** minimal lentivector with GFP in place of *gag*, SFFV promoter driving expression of *tat*, *rev*, or both, and repaired U3 in the 3’ LTR; the latter restores 5’-LTR-directed transcription to the provirus as a result of the reverse transcription strand-transfer reactions. When an essential viral component was disrupted within HIV-1-GFP, the factor in question was provided *in trans*, either during assembly in transfected HEK293 cells, or within transduced DCs, as appropriate (see Methods). Shown are blood donor data representative of n=6 (**a, b, e, f**), n=12 (**c, g, h**), n=8 (**d**). To determine significance, the MFI of individual flow cytometry samples was calculated as fold-change versus control. When data from each donor replicate within a experiment was combined, the difference in MFI for all experimental vs control conditions was significant in all cases, p<0.0001; one-way ANOVA, Dunnett’s post-test against HIV-1-GFP for (**a**, **c**, **d, h**) or lentivector control for (**e, f**). qRT-PCR data in are mean+/SD (p < 0.0001; two-way ANOVA, Dunnett’s post-test).

HIV-1 virion RNA and newly synthesized viral cDNA are reported to be detected by RIG-I and by cGAS, respectively (Berg et al., 2012; Gao et al., 2013). Signal transduction downstream of both sensors requires TBK1 and IRF3. The TBK1 inhibitor BX795 (Table 2) blocked DC maturation in response to cGAMP but had no effect on maturation after HIV-1-GFP transduction (Fig. 2c). Moreover, IRF3 knockdown (Table 1) (Pertel et al., 2011) suppressed activation of CD86 or ISG15 in response to cGAMP, but not in response to HIV-1 transduction (Fig. 2c). Similarly, no effect on HIV-1-induced DC maturation was observed with knockdown of IRF1, 5, 7, or 9, or of STAT1 or 2, or TAK1 (Table 1) (Pertel et al., 2011), or of pharmacologic inhibition of CypA, PKR, c-Raf, IkBa, NF-kB, MEK1+2, p38, JNK, Caspase 1, pan-Caspases, ASK1, eIF2a, TBK1, IKKe, TAK1, or NLRP3 (Table 2). Under the conditions used here, then, DC maturation required reverse transcription and integration but was independent of most well-characterized innate immune signaling pathways.

Completion of the HIV-1 integration reaction requires cellular DNA repair enzymes (Craigie and Bushman, 2012). That DCs did not mature in response to transduction with minimal lentivectors (Fig. 1b, i, j) indicates that activation of the DNA repair process is not sufficient, and that transcription from the HIV-1-GFP provirus must be necessary for maturation. Indeed, *gag* expression from an integrated vector has been reported to be necessary for DC maturation (Manel et al., 2010). To determine if any individual HIV-1 proteins were sufficient to mature DCs, a minimal lentivector was used to express codon optimized versions of each of the open reading frames possessed by HIV-1-GFP (Fig. 2d, Table 1). Among these vectors was a *gag*-expression vector that produced as much p24 protein as did HIV-1-GFP (Fig. 2d). None of these vectors matured DCs (Fig. 2d).

HIV-1-GFP was then mutated to determine if any protein coding sequences were necessary for DC maturation. For these and any subsequent experiments in which an essential viral component was disrupted within HIV-1-GFP, the factor in question was provided *in trans*, either during assembly in transfected HEK293 cells, or within transduced DCs, as appropriate (see Methods). Mutations that disrupted both *gag* and *pol*, either a double frameshift in *gag*, or a mutant in which the first 14 ATGs in *gag* were mutated, abolished synthesis of CA (p24) yet retained full maturation activity (Fig. 2e, Table 1). Deletion mutations encompassing *gag*/*pol*, *vif*/*vpr*, *vpu*/*env*, or *nef*/U3-LTR, each designed so as to leave *cis*-acting RNA elements intact, all matured DCs (Fig. 2f and Table 1). These results indicate that these HIV-1-GFP RNA sequences, as well as the proteins that they encode, were not required for DC maturation.

Tat and Rev coding sequences were individually disrupted by combining start codon point mutations with nonsense codons that were silent with respect to overlapping reading frames (Table 1). Neither Δ*tat* nor Δ*rev* matured DCs upon transduction (Fig. 2g). However, DCs matured upon co-transduction of Δ*tat* and Δ*rev*, or when minimal lentivectors expressing codon-optimized Tat and Rev were co-transduced *in trans* (Fig. 2g). These results indicate that the maturation defect with the individual vectors was due to disruption of Tat and Rev function, and not due to a *cis*-acting defect of the mutant RNA.

The minimal 3-part lentivector expressed GFP from a heterologous promoter and had a deletion mutation encompassing the essential, *cis*-acting TATA box and enhancer elements (Zufferey et al., 1998), as well as in the *trans*-acting *tat* and *rev*, that inactivated the promoter in the proviral 5’ LTR (Fig. 1a). To test the importance of LTR-driven transcription for DC maturation by the HIV-1 provirus, the HIV-1 LTR was restored in the minimal vector (Fig. 2h and Table 1); in addition, GFP was inserted in place of *gag* as a marker for LTR expression, and the heterologous promoter was used to drive *tat*, *rev*, or both genes separated by P2A coding sequence (Fig. 2h). None of the LTR-driven, minimal vectors matured DCs (Fig. 2h).

To determine if *tat* was necessary for DC maturation, *tat* and TAR were mutated in HIV-1-GFP and the LTR promoter was modified to be tetracycline-inducible, as previously described (Das and Berkhout, 2016) (Tet-HIV-1 in Fig. 3a and Table 1). The doxycycline-dependent reverse transactivator (*rtTA*) was delivered *in trans* by lentivector. In the presence of doxycycline (Table 2), Tet-HIV-1 and rtTA matured DCs when given in combination, but neither vector matured DCs when given in isolation (Fig. 3a). Additionally, the magnitude of cell surface CD86 was dependent on the doxycycline concentration, indicating that maturation was dependent on the level of HIV-1 transcription (Fig. 3b). These results demonstrated that *tat* was not required for maturation, so long as the provirus was expressed.

**Figure 3.**
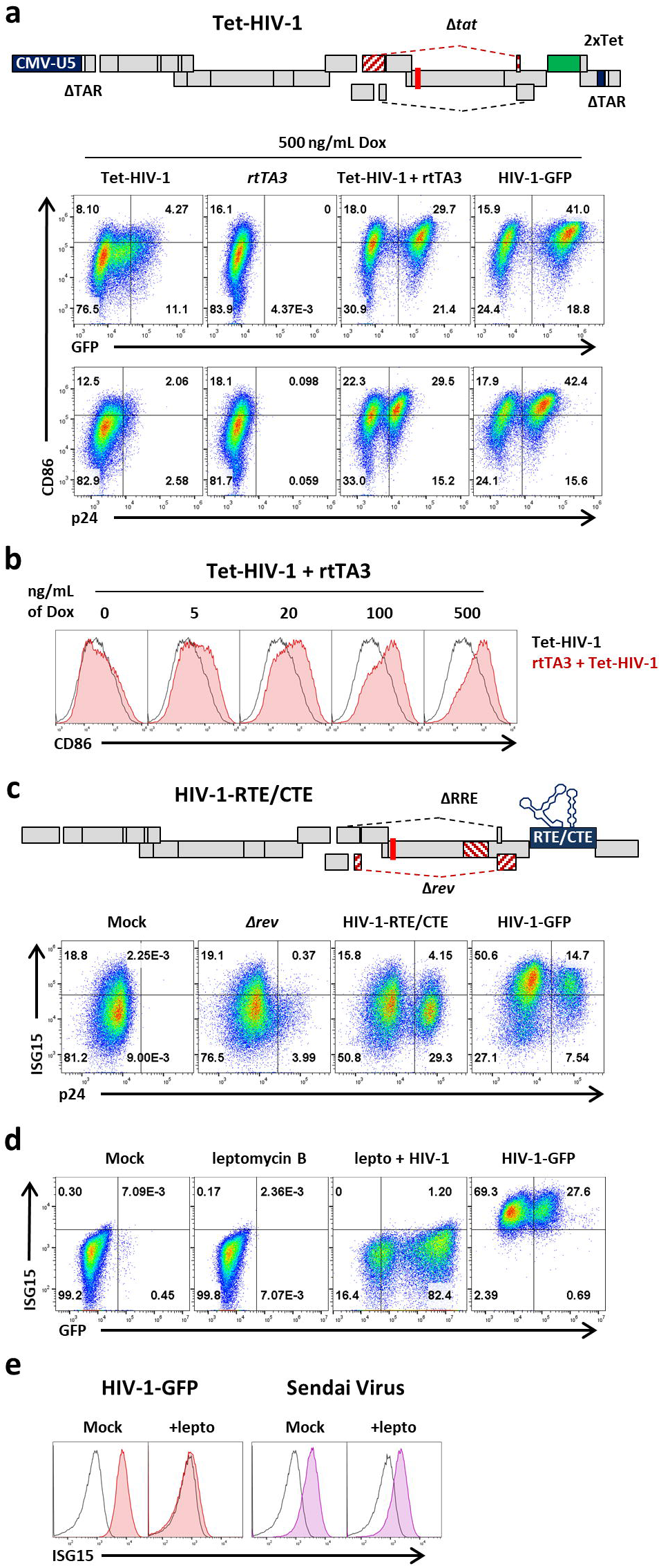
Rev-mediated RNA export is necessary for DC maturation but Tat is dispensable. **a**, Optimized 2xTet operator (Das and Berkhout, 2016) was cloned into the 3’LTR of HIV-1-GFPΔ*tat* to generate Tet-HIV-1; the strand-transfer reactions that occur during reverse transcription generate a Tet-regulated 5’-LTR in the provirus. DCs transduced with Tet-HIV-1, rtTA3, or both, were treated for 3 d with 500 ng/mL doxycycline and assayed by flow cytometry for p24, GFP, and CD86. **b**, DCs co-transduced with Tet-HIV-1 and rtTA3 were treated with increasing concentrations of doxycycline. **c**, To generate HIV-1-RTE/CTE, the RTEm26CTE element (Smulevitch et al., 2006) was cloned in place of *nef* in HIV-1-GFPΔ*rev*/ΔRRE. DCs were transduced with the indicated vectors and assessed for p24 and ISG15 by flow cytometry. **d**, DCs were treated with 25 nM leptomycin B, transduced with HIV-1-GFP, and assessed for GFP and ISG15 by flow cytometry. **e**, DCs were treated with 25 nM leptomycin B, transduced with HIV-1-GFP or infected with Sendai virus (SeV), and assessed for ISG15 by flow cytometry. Shown are blood donor data representative of n=10 (**a, c**), n=4 (**b**), n=6 (**d, e**). To determine significance, the MFI of individual flow cytometry samples was calculated as fold-change versus control. When data from each donor replicate within a experiment was combined, the difference in MFI for all experimental vs control conditions was significant in all cases, p<0.0001; one-way ANOVA, Dunnett’s post-test against dox negative control for (**a, b**) or HIV-1-GFP for (**c, d, e**).

To ascertain whether *rev* was necessary for DC maturation, the RTE from a murine intracisternal A-particle retroelement (IAP), and the CTE from SRV-1, were inserted in place of *nef* (HIV-RTE/CTE in Fig. 3c and Table 1) (Smulevitch et al., 2006). Each of these elements utilizes the NXF1 nuclear RNA export pathway, thereby bypassing the need for CRM1 and *rev (Fornerod et al., 1997)*. p24 levels with this construct were similar to those of HIV-1-GFP, indicating that unspliced RNA was exported from the nucleus at least as well as with Rev (Fig. 3c). Nonetheless, the HIV-RTE/CTE vector did not mature DCs (Fig. 3c), indicating that maturation was dependent upon *rev* and CRM1-mediated RNA export. Consistent with this conclusion, the CRM1 inhibitor leptomycin B (Table 2) abrogated DC maturation by HIV-1-GFP (Fig. 3d). In contrast, leptomycin B had no effect on DC maturation in response to Sendai virus infection (Fig. 3e). ISG15 was used to monitor maturation in these experiments since, as previously reported for DCs, leptomycin B altered background levels of CD86 (Chemnitz et al., 2010).

To determine if innate immune detection of HIV-1 was unique to DCs, monocyte-derived macrophages and CD4^+^ T cells were examined. In response to transduction with HIV-1-GFP, macrophages upregulated CD86, ISG15, and HLA-DR, and CD4^+^ T cells upregulated MX1, IFIT1, and HLA-DR (Fig. 4a). DCs, macrophages, and CD4^+^ T cells were then transduced side-by-side with mutant constructs to determine if the mechanism of innate immune activation was similar to that in DCs. As with DCs, HIV-1-GFP bearing the Δ*gag/pol* deletion activated macrophages and CD4^+^ T cells (Fig. 4b). Also in agreement with the DC results, neither the minimal lentivector, nor HIV-1-GFP bearing mutations in *integrase*, *tat*, or *rev*, matured any of the three cell types (Fig. 4b). CD4^+^ T cells were infected with either macrophage-tropic or T cell-tropic HIV-1 to determine whether replication-competent HIV-1 was similarly capable of innate immune activation in these cells, in the absence of VSV G. As with DCs, innate immune activation, as detected by MX1 and ISG15 upregulation, was observed in cells productively infected with HIV-1, but not with minimal lentivector (Fig. 4c). Finally, to test the effect of HIV-1 proviral RNA on non-activated T cells, CD4^+^ T cells were co-transduced with Tet-HIV-1 and the rtTA3 vector, and cultured for 9 days in the absence of stimulation. Upon doxycycline treatment, T cells expressed GFP and MX1 (Fig. 4d). As in DCs, dose-dependent activation was observed with doxycycline (Fig. 4d). These data indicate that innate immune activation by HIV-1, in all three cell types, requires integration, transcription, and Rev-dependent, HIV-1 intron-containing RNA.

**Figure 4.**
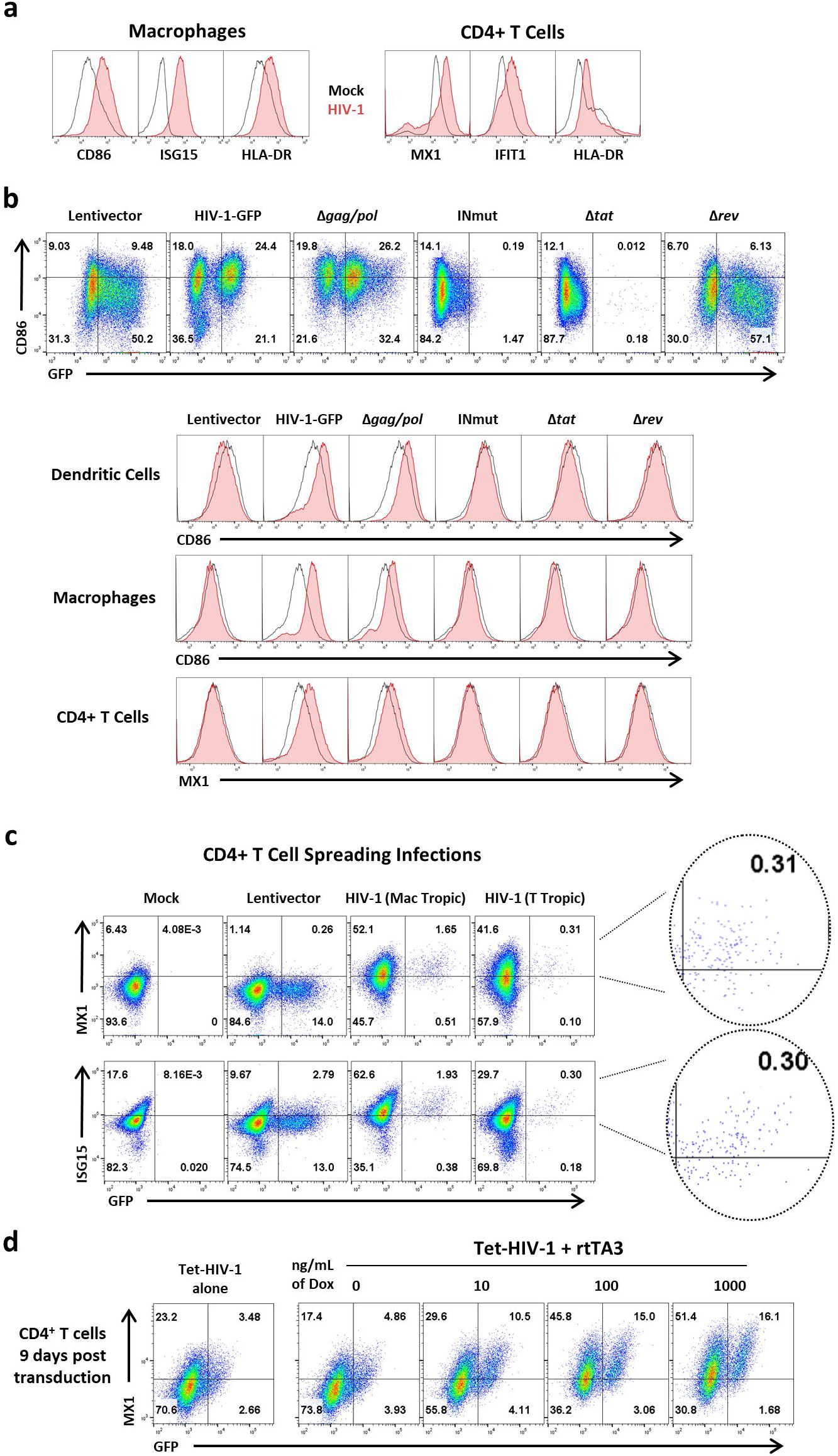
Innate immune activation in macrophages and CD4^+^ T cells by HIV-1 proviral transcription. **a**, Macrophages and CD4^+^ T cells were transduced with HIV-1-GFP and assayed 3 days later for the indicated activation markers. **b**, DCs, macrophages, and CD4^+^ T cells were challenged with HIV-1-GFP or the indicated mutants. When an essential viral component was disrupted within HIV-1-GFP, the factor in question was provided *in trans* during assembly in transfected HEK293 cells, as appropriate (see Methods). The upper panel shows flow cytometry of the DCs for GFP and CD86. The histograms show CD86 for DCs and macrophages or MX1 for CD4^+^ T cells. **c**, 12 day spreading infections on CD4+ T cells with either macrophage-tropic or T cell-tropic, replication-competent HIV-1. **d,** CD4^+^ T cells were stimulated for 3 days with PHA and IL2, and transduced with Tet-HIV-1 and rtTA3. Cells were then cultured without stimulation for 9 days. Doxycycline was then added at the indicated concentrations. Cells were assayed for GFP and MX1 3 days later. Shown are blood donor data representative of n=6 (**a**), n=4 (**b, c, d**). To determine significance, the MFI of individual flow cytometry samples was calculated as fold-change versus control. The exception being (**c**) where the MFI of only GFP+ cells was compared. When data from each donor replicate within a experiment was combined, the difference in MFI for all experimental vs control conditions was significant in all cases, p<0.0001; one-way ANOVA, Dunnett’s post-test against lentivector control for (**a, b, c**) or dox control for (**d**).

## DISCUSSION

The HIV-1 LTR generates a single primary transcript that gives rise to over 100 alternatively spliced RNAs (Ocwieja et al., 2012). The full-length, unspliced, intron-bearing transcript acts as viral genomic RNA in the virion and as mRNA for essential *gag*- and *pol*-encoded proteins. Expression of the unspliced transcript requires specialized viral and cellular machinery, HIV-1 Rev and CRM1 (Fornerod et al., 1997), in order to escape from the spliceosome. Results here indicate that unspliced or partially spliced HIV-1 RNA is detected by human cells as a danger signal, as has been reported for inefficiently spliced mRNAs from transposable elements in distantly related eukaryotes (Dumesic et al., 2013). Transposable elements are mutagenic to the host genome and it stands to reason that molecular features such as transcripts bearing multiple, inefficient splice signals characteristic of retrotransposons, would activate innate immune signaling pathways.

HIV-1 genomic RNA contains extensive secondary and higher order structures that could be detected by innate immune sensors. Our knockdown of IRF3 and inhibition of TBK1, both required for signal transduction of the RNA sensors RIG-I, MDA5, and TLR3, did not impede HIV-1 maturation of DCs (Figure 2c). Furthermore, we suppressed an extensive list of innate signaling pathways and sensors including knockdowns of IRF’s 1, 5, 7, and 9, STAT’s 1 and 2, or of TAK1, as well as pharmacologic inhibition of CypA, CRM1, PKR, c-Raf, IkBa, NF-kB, MEK1+2, p38, JNK, Caspase 1, pan-Caspases, ASK1, eIF2a, IKKe, TAK1, or NLRP3 (Table 2). None of these perturbations had any effect on limiting innate immune activation by HIV-1, suggesting requirement for an alternative detection mechanism. Such mechanisms might include uncharacterized RNA sensors, direct detection of stalled splicing machinery, or overload of the CRM1 nuclear export pathway itself.

The replication competent HIV-1 reservoir in memory CD4^+^ T cells has a 44 wk half-life and thus patients must take antiviral medication for life (Crooks et al., 2015). Long-lived, replication competent HIV-1 reservoirs in other cell types have not been clearly demonstrated, but these may also contribute to the HIV-1 reservoir (Kandathil et al., 2016). The common genetic determinants in HIV-1 for maturation of CD4^+^ T cells, macrophages, and DCs suggests that HIV-1 is detected by a mechanism that is conserved across cell types, and that this mechanism would be active in any cell type that possesses a transcriptionally active provirus. Data here show that proviruses need not be replication competent to contribute to inflammation. Rather, HIV-1 transcription and export of unspliced RNA, regardless of replication competence, is sufficient to induce immune activation. Consistent with our findings, T cell activation correlates directly with the level of cell-associated HIV-1 RNA in patients receiving antiretroviral therapy (El-Diwany et al., 2017). Furthermore, our data suggests that new drugs that block HIV-1 transcription, Tat-mediated transcriptional elongation, or Rev-mediated preservation of unspliced transcripts (Mousseau et al., 2015), would limit inflammation, and offer an important addition to the current anti-HIV-1 drug armamentarium.

## METHODS

### Data reporting

No statistical methods were used to predetermine sample size. The experiments were not randomized. The investigators were not blinded to allocation during experiments and outcome assessment.

### Plasmids

The plasmids used here were either previously described or generated using standard cloning methods (Pertel et al., 2011). The full list of plasmids used here, along with their purpose and characteristics, is provided in Table 1. All plasmid DNAs with complete nucleotide sequence files are available at www.addgene.com.

### Cell culture

Cells were cultured at 37°C in 5% CO_2_ humidified incubators and monitored for mycoplasma contamination using the Lonza Mycoplasma Detection kit by Lonza (LT07-318). HEK293 cells (ATCC) were used for viral production and were maintained in DMEM supplemented with 10% FBS, 20 mM L-glutamine (ThermoFisher), 25 mM HEPES pH 7.2 (SigmaAldrich), 1 mM sodium pyruvate (ThermoFisher), and 1x MEM non-essential amino acids (ThermoFisher). Cytokine conditioned media was produced from HEK293 cells stably transduced with pAIP-hGMCSF-co (Addgene #74168), pAIP-hIL4-co (Addgene #74169), or pAIP-hIL2 (Addgene #90513), as previously described (Pertel et al., 2011).

Leukopaks were obtained from anonymous, healthy, blood bank donors (New York Biologics). As per NIH guidelines (http://grants.nih.gov/grants/policy/hs/faqs_aps_definitions.htm), experiments with these cells were declared non-human subjects research by the UMMS IRB. PBMCs were isolated from leukopaks by gradient centrifugation on Histopaque 1077 (Sigma-Aldrich). CD14^+^ mononuclear cells were isolated via positive selection using anti-CD14 antibody microbeads (Miltenyi). Enrichment for CD14^+^ cells was routinely>X98%.

To generate DCs or macrophages, CD14^+^ cells were plated at a density of 1 to 2 × 10^6^ cells/ml in RPMI-1640 supplemented with 5% heat inactivated human AB^+^ serum (Omega Scientific, Tarzana, CA), 20 mM L-glutamine, 25 mM HEPES pH 7.2, 1 mM sodium pyruvate, and 1x MEM non-essential amino acids (RPMI-HS complete) in the presence of cytokines that promote differentiation. DCs were generated by culturing monocytes for 6 days in the presence of 1:100 cytokine-conditioned media containing human GM-CSF and human IL-4. DC preparations were consistently >99% DC-SIGN^high^, CD11c^high^, and CD14^low^ by flow cytometry. Macrophages were generated by culturing for 7 days with GM-CSF conditioned media in the absence of IL-4, and were routinely >99% CD11b. CD4^+^ T cells were isolated from PBMCs that had been depleted of CD14^+^ cells, as above, using anti-CD4 microbeads (Miltenyi), and were >99% CD4^+^. CD4^+^ T cells were then cultured in RPMI-1640 supplemented with 10% heat inactivated FBS, 20 mM L-glutamine, 25 mM HEPES pH 7.2, 1 mM sodium pyruvate, 1x MEM non-essential amino acids (RPMI-FBS complete), and 1:2000 IL-2 conditioned media. Cells from particular donors were excluded from experiments if percent enrichment deviated more than 5% from the numbers mentioned above, or if there was no increase in activation markers in response to control stimuli (LPS, Sendai virus, and wild-type HIV-1-GFP).

### HIV-1 vector production

HEK293E cells were seeded at 75% confluency in 6-well plates and transfected with 6.25 uL Transit LT1 lipid reagent (Mirus) in 250 μL Opti-MEM (Gibco) with 2.25 µg total plasmid DNA. 2-part HIV-1 vectors based on HIV-1-GFP (He et al., 1997) and described in detail in Table 1 were transfected at a 7:1 ratio in terms of µgs of HIV-1 plasmid DNA to pMD2.G VSV G expression plasmid DNA (Pertel et al., 2011). 3-part lentivectors were produced by transfection of the lentivector genome, psPAX2 GagPol vector, and pMD2.G, at a DNA ratio of 4:3:1. These also include 2-part HIV-1-GFP constructs that are mutated in such a way as to prevent GagPol, Tat, or Rev production. As these would be defective for viral production, psPAX2 was included in the transfections at the same 4:3:1 ratio. VPX-bearing SIV-VLPs were produced by transfection at a 7:1 plasmid ratio of SIV3+ to pMD2.G (Pertel et al., 2011). 12 hrs after transfection, media was changed to the specific media for the cells that were to be transduced. Viral supernatant was harvested 2 days later, filtered through a 0.45 µm filter, and stored at 4°C.

Virions in the transfection supernatant were quantified by a PCR-based assay for reverse transcriptase activity (Pertel et al., 2011). 5 μl transfection supernatant were lysed in 5 μL 0.25% Triton X-100, 50 mM KCl, 100 mM Tris-HCl pH 7.4, and 0.4 U/μl RNase inhibitor (RiboLock, ThermoFisher). Viral lysate was then diluted 1:100 in a buffer of 5 mM (NH_4_)_2_SO_4_, 20 mM KCl, and 20 mM Tris–HCl pH 8.3. 10 μL was then added to a single-step, RT PCR assay with 35 nM MS2 RNA (IDT) as template, 500 nM of each primer (5’-TCCTGCTCAACTTCCTGTCGAG-3’ and 5’-CACAGGTCAAACCTCCTAGGAATG-3’), and hot-start Taq (Promega) in a buffer of 20 mM Tris-Cl pH 8.3, 5 mM (NH_4_)_2_SO_4_, 20 mM KCl, 5 mM MgCl_2_, 0.1 mg/ml BSA,

1/20,000 SYBR Green I (Invitrogen), and 200 μM dNTPs. The RT-PCR reaction was carried out in a Biorad CFX96 cycler with the following parameters: 42°C 20 min, 95°C 2 min, and 40 cycles [95°C for 5 s, 60°C 5 s, 72°C for 15 s and acquisition at 80°C for 5 s]. 2 part vectors typically yielded 10^7^ RT units/µL, and 3 part vector transfections yielded 10^6^ RT units/µL.

### Transductions

10^6^ DCs/mL, or 5 × 10^5^ macrophages/ml, were plated into RPMI-HS complete with Vpx^+^ SIV-VLP transfection supernatant added at a dilution of 1:6. After 2 hrs, 10^8^ RT units of viral vector was added. In some cases, drugs were added to the culture media as specified in Table 2. In most cases, transduced DC were harvested for analysis 3 days following challenge. For gene knockdown or for expression of factors *in trans*, 2 × 10^6^ CD14^+^ monocytes/mL were transduced directly following magnetic bead isolation with 1:6 volume of SIV-VLPs and 1:6 volume of vector. When drug selection was required, 4 µg/mL puromycin was added 3 days after monocyte transduction and cells were selected for 3 days. SIV-VLPs were re-administered in all cases with HIV-1 or lentivector challenge. For DCs in Tet-HIV-1 experiments, fresh monocytes were SIV-VLP treated and co-transduced with rtTA3 and Tet-HIV-1. DCs were harvested 6 days later and treated with indicated doxycycline concentrations.

For deoxynucleoside-assisted transductions, DCs were plated at 10^6^ DCs/mL and treated with 2mM of combined deoxynucleosides for 2 hrs before transduction with HIV-1. Deoxynucleosides were purchased from Sigma-Aldrich (2′deoxyguanosine monohydrate, cat# D0901; thymidine, cat# T1895; 2′deoxyadenosine monohydrate, cat# D8668; 2′deoxycytidine hydrochloride, cat# D0776). A 100 mM stock solution was prepared by dissolving each of the four nucleotides at 100 mM in RPMI 1640 by heating the medium at 80°C for 15 min.

CD4^+^ T cells were stimulated in RPMI-FBS complete with 1:2000 hIL-2 conditioned media and 5 μg/mL PHA-P. After 3 days, T cells were replated at 10^6^ cells/mL in RPMI-FBS complete with hIL-2. Cells were transduced with 10^8^ RT units of viral vector per 10^6^ cells and assayed 3 days later. T cells were co-transduced with rtTA3 and Tet-HIV-1 every day for 3 days after PHA stimulation. Cells were then replated in RPMI-FBS complete with hIL-2. Transduced T cells were cultured for 9 days with fresh media added at day 5. After 9 days, doxycycline was added at the indicated concentrations and assayed 3 days later.

### Non-HIV-1 Challenge Viruses

Sendai Virus Cantell Strain was purchased from Charles River Laboratories. Infections were performed with 200 HA units/ml on DCs for 3 days before assay by flow cytometry.

### Spreading Infections

DCs were plated at 10^6^ DCs/mL, in RPMI-HS complete media, with or without Vpx^+^ SIV-VLP transfection supernatant added at a dilution of 1:6. After 2 hrs, 10^8^ RT units of HEK-293 transfection supernatant of either NL4-3-GFP with JRCSF env (T tropic) or NL4-3-GFP with JRFL env (mac tropic) was added. Every 3 days (for a total of 12 days) samples were harvested for detection of viral RT activity in supernatant and flow cytometry assessment.

CD4^+^ T cells were stimulated in RPMI-FBS complete with 1:2000 hIL-2 conditioned media and 5 μg/mL PHA-P. After 3 days, T cells were replated at 10^6^ cells/mL in RPMI-FBS complete with hIL-2 and transduced with 10^8^ RT units of NL4-3-GFP with JRFL (mac tropic) or JRCSF (T tropic) env. Cells were harvested every 3 days (for a total of 12 days) and assayed for infectivity and activation via flow cytometry.

### Cytokine analysis

Supernatants from DCs were collected 3 days following transduction with HIV-1-GFP or minimal lentivector. Supernatant was spun at 500 × g for 5 mins and filtered through a 0.45 μm filter. Multiplex soluble protein analysis was carried out by Eve Technologies (Calgary, AB, Canada).

### qRT-PCR

Total RNA was isolated from 5 × 10^5^ DCs using RNeasy Plus Mini (Qiagen) with Turbo DNase (ThermoFisher) treatment between washes. First-strand synthesis used Superscript III Vilo Master mix (Invitrogen) with random hexamers. qPCR was performed in 20 μL using 1× TaqMan Gene Expression Master Mix (Applied Biosystems), 1 μL cDNA, and 1 μL TaqMan Gene Expression Assays (ThermoFisher) specified in Table 3. Amplification was on a CFX96 Real Time Thermal Cycler (Bio-Rad) using the following program: 95°C for 10 min [45 cycles of 95°C for 15 s and 60°C for 60 s]. Housekeeping gene OAZ1 was used as control (Pertel et al., 2011).

**Table 3.** qRT-PCR probes.

| **RNA Target** | **Taqman probe ID#** |
| --- | --- |
| CXCL10 | Hs00171042_m1 |
| IFNB1 | Hs01077958_s1 |
| IL15 | Hs01003716_m1 |
| OAZ1 | Hs00427923_m1 |
| ACTB | Hs01060665_g1 |

### Flow cytometry

10^5^ cells were surface stained in FACS buffer (PBS, 2% FBS, 0.1% Sodium Azide), using the antibodies in Table 4. Cells were then fixed in a 1:4 dilution of BD Fixation Buffer and assayed on a BD C6 Accuri. BD Biosciences Fixation and Permeabilization buffers were utilized for intracellular staining. Data was analyzed in FlowJo.

**Table 4.** Antibodies used in this study.

| **Target antigen** | **Clone** | **Fluorophore** | **Source** |
| --- | --- | --- | --- |
| CD80 | 2D10 | PE, APC | Biolegend |
| CD86 | IT2.2 | PE, APC | Biolegend |
| CD40 | HB14 | APC | Biolegend |
| HLA-DR | L249 | PerCP, APC | Biolegend |
| CD83 | HB15e | FITC, APC | Biolegend |
| CCR7 | G043H7 | PE | Biolegend |
| CD141 | M80 | PE | Biolegend |
| ISG15 | #851701 | PE, APC | R&D Systems |
| MX1 | EPR19967 | N/A | Abcam |
| IFIT | OTI3G8 | N/A | Abcam |
| p24 | KC57 | FITC, PE | BeckmanCoulter |
| CD1a | HI149 | FITC, PerCP | Biolegend |
| CD1c | L161 | PE | Biolegend |
| CD14 | HCD14 | FITC, Pe-Cy7 | Biolegend |
| CD11b | ICRF44 | PE | Biolegend |
| CD11c | 3.9 | PE, PE-Cy7, APC | Biolegend |
| CD209 (DC-SIGN) | 9E9A8 | APC | Biolegend |
| CXCL10 | JO34D6 | PE | Biolegend |
| HLA-ABC | W6/32 | FITC | Biolegend |
| CD3 | OKT3 | FITC, APC, BV650 | Biolegend |
| CD4 | OKT4 | FITC, APC, Alexa700 | Biolegend |
| Mouse-IgG | Poly4053 | PE, APC | Biolegend |
| Rabbit-IgG | Poly4064 | PE, AlexaFluor647 | Biolegend |

### Sampling

All individual experiments were performed with biological duplicates, using cells isolated from two different blood donors. Flow cytometry plots in the figures show representative data taken from experiments performed with cells from the number of donors indicated in the figure legends.

### Statistical Analysis

Experimental n values and information regarding specific statistical tests can be found in the figure legends. The mean fluorescence intensity for all live cells analyzed under a given condition was calculated as fold-change to negative control/mock. The exception to this methodology was in Figures 1g and 4c where the percent infected cells was too low to use MFI for the bulk population; in these cases MFI was determined for the subset of cells within the GFP+ gate. Significance of flow cytometry data was determined via one-way ANOVA. A Dunnett’s post-test for multiple comparisons was applied, where MFI fold change was compared to either mock treatment or positive treatment depending on the experimental question. qRT-PCR and luminex data was analyzed via two-way Anova, with Dunnett’s post-test comparing all samples to mock. All ANOVAs were performed using PRISM 7.02 software (GraphPad Software, La Jolla, CA).

## Data availability

The plasmids described in Table 1, along with their complete nucleotide sequences, are available at www.addgene.com.

## ACKNOWLEDGEMENTS

We thank the 256 anonymous blood donors who contributed leukocytes to this project, and Ben Berkhout, Abraham Brass, Barbara Felber, Massimo Pizzato, and Didier Trono for advice and reagents. This work was supported by NIH Grants 5R01AI111809, 5DP1DA034990, and 1R01AI117839, to J.L. The plasmids described in Table 1, along with their complete nucleotide sequences, are available at www.addgene.com.

## REFERENCES

Berg, R.K., Melchjorsen, J., Rintahaka, J., Diget, E., Søby, S., Horan, K.A., Gorelick, R.J., Matikainen, S., Larsen, C.S., Ostergaard, L., et al. (2012). Genomic HIV RNA induces innate immune responses through RIG-I-dependent sensing of secondary-structured RNA. PLoS One 7, e29291.

Brenchley, J.M., Price, D.A., Schacker, T.W., Asher, T.E., Silvestri, G., Rao, S., Kazzaz, Z., Bornstein, E., Lambotte, O., Altmann, D., et al. (2006). Microbial translocation is a cause of systemic immune activation in chronic HIV infection. Nat. Med. 12, 1365–1371.

Bruner, K.M., Murray, A.J., Pollack, R.A., Soliman, M.G., Laskey, S.B., Capoferri, A.A., Lai, J., Strain, M.C., Lada, S.M., Hoh, R., et al. (2016). Defective proviruses rapidly accumulate during acute HIV-1 infection. Nat. Med. 22, 1043–1049.

Chemnitz, J., Turza, N., Hauber, I., Steinkasserer, A., and Hauber, J. (2010). The karyopherin CRM1 is required for dendritic cell maturation. Immunobiology 215, 370–379.

Craigie, R., and Bushman, F.D. (2012). HIV DNA integration. Cold Spring Harb. Perspect. Med. 2, a006890.

Crooks, A.M., Bateson, R., Cope, A.B., Dahl, N.P., Griggs, M.K., Kuruc, J.D., Gay, C.L., Eron, J.J., Margolis, D.M., Bosch, R.J., et al. (2015). Precise quantitation of the latent HIV-1 reservoir: implications for eradication strategies. J. Infect. Dis. 212, 1361–1365.

Das, A.T., and Berkhout, B. (2016). Conditionally replicating HIV and SIV variants. Virus Res. 216, 66–75.

Davey, R.T., Jr, Bhat, N., Yoder, C., Chun, T.W., Metcalf, J.A., Dewar, R., Natarajan, V., Lempicki, R.A., Adelsberger, J.W., Miller, K.D., et al. (1999). HIV-1 and T cell dynamics after interruption of highly active antiretroviral therapy (HAART) in patients with a history of sustained viral suppression. Proc. Natl. Acad. Sci. U. S. A. 96, 15109–15114.

De Iaco, A., and Luban, J. (2011). Inhibition of HIV-1 infection by TNPO3 depletion is determined by capsid and detectable after viral cDNA enters the nucleus. Retrovirology 8, 98.

Deymier, M.J., Ende, Z., Fenton-May, A.E., Dilernia, D.A., Kilembe, W., Allen, S.A., Borrow, P., and Hunter, E. (2015). Heterosexual Transmission of Subtype C HIV-1 Selects Consensus-Like Variants without Increased Replicative Capacity or Interferon-α Resistance. PLoS Pathog. 11, e1005154.

Dumesic, P.A., Natarajan, P., Chen, C., Drinnenberg, I.A., Schiller, B.J., Thompson, J., Moresco, J.J., Yates, J.R., 3rd, Bartel, D.P., and Madhani, H.D. (2013). Stalled spliceosomes are a signal for RNAi-mediated genome defense. Cell 152, 957–968.

El-Diwany, R., Breitwieser, F.P., Soliman, M., Skaist, A.M., Srikrishna, G., Blankson, J.N., Ray, S.C., Wheelan, S.J., Thomas, D.L., and Balagopal, A. (2017). Intracellular HIV-1 RNA and CD4+ T-cell activation in patients starting antiretrovirals. AIDS 31, 1405–1414.

Engelman, A.N., and Singh, P.K. (2018). Cellular and molecular mechanisms of HIV-1 integration targeting. Cell. Mol. Life Sci.

Fornerod, M., Ohno, M., Yoshida, M., and Mattaj, I.W. (1997). CRM1 is an export receptor for leucine-rich nuclear export signals. Cell 90, 1051–1060.

Freiberg, M.S., Chang, C.-C.H., Kuller, L.H., Skanderson, M., Lowy, E., Kraemer, K.L., Butt, A.A., Bidwell Goetz, M., Leaf, D., Oursler, K.A., et al. (2013). HIV infection and the risk of acute myocardial infarction. JAMA Intern. Med. 173, 614–622.

Gao, D., Wu, J., Wu, Y.-T., Du, F., Aroh, C., Yan, N., Sun, L., and Chen, Z.J. (2013). Cyclic GMP-AMP synthase is an innate immune sensor of HIV and other retroviruses. Science 341, 903–906.

Goujon, C., Jarrosson-Wuillème, L., Bernaud, J., Rigal, D., Darlix, J.-L., and Cimarelli, A. (2006). With a little help from a friend: increasing HIV transduction of monocyte-derived dendritic cells with virion-like particles of SIV(MAC). Gene Ther. 13, 991–994.

Granelli-Piperno, A., Delgado, E., Finkel, V., Paxton, W., and Steinman, R.M. (1998). Immature Dendritic Cells Selectively Replicate Macrophagetropic (M-Tropic) Human Immunodeficiency Virus Type 1, while Mature Cells Efficiently Transmit both M- and T-Tropic Virus to T Cells. J. Virol. 72, 2733–2737.

Günthard, H.F., Saag, M.S., Benson, C.A., del Rio, C., Eron, J.J., Gallant, J.E., Hoy, J.F., Mugavero, M.J., Sax, P.E., Thompson, M.A., et al. (2016). Antiretroviral Drugs for Treatment and Prevention of HIV Infection in Adults: 2016 Recommendations of the International Antiviral Society-USA Panel. JAMA 316, 191–210.

He, J., Chen, Y., Farzan, M., Choe, H., Ohagen, A., Gartner, S., Busciglio, J., Yang, X., Hofmann, W., Newman, W., et al. (1997). CCR3 and CCR5 are co-receptors for HIV-1 infection of microglia. Nature 385, 645–649.

Hunt, P.W., Martin, J.N., Sinclair, E., Epling, L., Teague, J., Jacobson, M.A., Tracy, R.P., Corey, L., and Deeks, S.G. (2011). Valganciclovir reduces T cell activation in HIV-infected individuals with incomplete CD4+ T cell recovery on antiretroviral therapy. J. Infect. Dis. 203, 1474–1483.

Jiang, A.-P., Jiang, J.-F., Wei, J.-F., Guo, M.-G., Qin, Y., Guo, Q.-Q., Ma, L., Liu, B.-C., Wang, X., Veazey, R.S., et al. (2015). Human Mucosal Mast Cells Capture HIV-1 and Mediate Viral trans-Infection of CD4+ T Cells. J. Virol. 90, 2928–2937.

Kandathil, A.J., Sugawara, S., and Balagopal, A. (2016). Are T cells the only HIV-1 reservoir? Retrovirology 13, 86.

Landau, N.R. (2014). The innate immune response to HIV-1: to sense or not to sense. DNA Cell Biol. 33, 271–274.

Manel, N., Hogstad, B., Wang, Y., Levy, D.E., Unutmaz, D., and Littman, D.R. (2010). A cryptic sensor for HIV-1 activates antiviral innate immunity in dendritic cells. Nature 467, 214–217.

Mousseau, G., Kessing, C.F., Fromentin, R., Trautmann, L., Chomont, N., and Valente, S.T. (2015). The Tat Inhibitor Didehydro-Cortistatin A Prevents HIV-1 Reactivation from Latency. MBio 6, e00465.

Ocwieja, K.E., Sherrill-Mix, S., Mukherjee, R., Custers-Allen, R., David, P., Brown, M., Wang, S., Link, D.R., Olson, J., Travers, K., et al. (2012). Dynamic regulation of HIV-1 mRNA populations analyzed by single-molecule enrichment and long-read sequencing. Nucleic Acids Res. 40, 10345–10355.

Parrish, N.F., Gao, F., Li, H., Giorgi, E.E., Barbian, H.J., Parrish, E.H., Zajic, L., Iyer, S.S., Decker, J.M., Kumar, A., et al. (2013). Phenotypic properties of transmitted founder HIV-1. Proc. Natl. Acad. Sci. U. S. A. 110, 6626–6633.

Pertel, T., Hausmann, S., Morger, D., Züger, S., Guerra, J., Lascano, J., Reinhard, C., Santoni, F.A., Uchil, P.D., Chatel, L., et al. (2011). TRIM5 is an innate immune sensor for the retrovirus capsid lattice. Nature 472, 361–365.

Rasaiyaah, J., Tan, C.P., Fletcher, A.J., Price, A.J., Blondeau, C., Hilditch, L., Jacques, D.A., Selwood, D.L., James, L.C., Noursadeghi, M., et al. (2013). HIV-1 evades innate immune recognition through specific cofactor recruitment. Nature 503, 402–405.

Reinhard, C., Bottinelli, D., Kim, B., and Luban, J. (2014). Vpx rescue of HIV-1 from the antiviral state in mature dendritic cells is independent of the intracellular deoxynucleotide concentration. Retrovirology 11, 12.

Salazar-Gonzalez, J.F., Salazar, M.G., Keele, B.F., Learn, G.H., Giorgi, E.E., Li, H., Decker, J.M., Wang, S., Baalwa, J., Kraus, M.H., et al. (2009). Genetic identity, biological phenotype, and evolutionary pathways of transmitted/founder viruses in acute and early HIV-1 infection. J. Exp. Med. 206, 1273–1289.

Siliciano, J.D., Kajdas, J., Finzi, D., Quinn, T.C., Chadwick, K., Margolick, J.B., Kovacs, C., Gange, S.J., and Siliciano, R.F. (2003). Long-term follow-up studies confirm the stability of the latent reservoir for HIV-1 in resting CD4+ T cells. Nat. Med. 9, 727–728.

Sinha, A., Ma, Y., Scherzer, R., Hur, S., Li, D., Ganz, P., Deeks, S.G., and Hsue, P.Y. (2016). Role of T-Cell Dysfunction, Inflammation, and Coagulation in Microvascular Disease in HIV. J. Am. Heart Assoc. 5, e004243.

Smulevitch, S., Bear, J., Alicea, C., Rosati, M., Jalah, R., Zolotukhin, A.S., von Gegerfelt, A., Michalowski, D., Moroni, C., Pavlakis, G.N., et al. (2006). RTE and CTE mRNA export elements synergistically increase expression of unstable, Rev-dependent HIV and SIV mRNAs. Retrovirology 3, 1–9.

Sousa, C.R. e. (2006). Dendritic cells in a mature age. Nat. Rev. Immunol. 6, 476–483.

Wiegand, A., Spindler, J., Hong, F.F., Shao, W., Cyktor, J.C., Cillo, A.R., Halvas, E.K., Coffin, J.M., Mellors, J.W., and Kearney, M.F. (2017). Single-cell analysis of HIV-1 transcriptional activity reveals expression of proviruses in expanded clones during ART. Proc. Natl. Acad. Sci. U. S. A. 114, E3659–E3668.

Zufferey, R., Dull, T., Mandel, R.J., Bukovsky, A., Quiroz, D., Naldini, L., and Trono, D. (1998). Self-inactivating lentivirus vector for safe and efficient in vivo gene delivery. J. Virol. 72, 9873–9880.

